# ontoFAST: An R package for interactive and semi-automatic annotation of characters with biological ontologies

**DOI:** 10.1101/2021.05.11.443562

**Authors:** Sergei Tarasov, István Mikó, Matthew Jon Yoder

## Abstract

1. The commonly used Entity-Quality (EQ) syntax provides rich semantics and high granularity for annotating phenotypes and characters using ontologies. However, EQ syntax might be time inefficient if this granularity is unnecessary for downstream analysis.
2. We present an R package ontoFAST that aid production of fast annotations of characters and character matrices with biological ontologies. Its interactive interface allows quick and convenient tagging of character statements with necessary ontology terms.
3. The annotations produced in ontoFAST can be exported in csv format for downstream analysis. Additinally, OntoFAST provides: (i) functions for constructing simple queries of characters against ontologies, and (ii) helper function for exporting and visualising complex ontological hierarchies and their relationships.
4. OntoFAST enhances data interoperability between various applications and support further integration of ontological and phylogenetic methods. Ontology tools are underrepresented in R environment and we hope that ontoFAST will stimulate their further development.

## 1 INTRODUCTION

During the past few decades a massive number of organismal phenotypes have been described by biologists for phylogenetic purposes in the form of character matrices and statements written in natural language (NL). Nowadays, ontologies are emerging as a fundamental technology for managing phenotypic data (Balhoff et al., 2010). An ontology is a computer-based representation of concepts and their logical relationships for a specific domain of knowledge (Deans et al., 2012, 2015; Balhoff et al., 2013). Ontological representation facilitates the conversion of NL into machine-parsable statements, thereby, providing new opportunities for computer-aided comparative phenomics and trait analysis (Deans et al., 2015; Dececchi et al., 2015; Burleigh et al., 2013; Tarasov, 2019).

The Entity-Quality (EQ) syntax, adopted by the Open Biological and Biomedical Ontologies (OBO) Consortium (Washington et al., 2009), and Phenoscape project (http://phenoscape.org), is the common convention for ontological representation of phenotypes (Balhoff et al., 2010; Gkoutos et al., 2005). EQ syntax bounds an entity, corresponding to a specific anatomical structure from an anatomy ontology with a quality term from the generic Phenotype and Trait Ontology [PATO, Mungall et al. (2010)]. The annotation of character matrices and phenotypes using EQ syntax is implemented in a comprehensive Java-based application Phenex (Balhoff et al., 2010), and an earlier software Phenote, designed for annotating mutant phenotype in model organisms (Washington et al., 2009). The flexibility of the EQ approach provides rich semantics for describing nearly any organismal phenotype at the very high level of granularity (Dahdul et al., 2018, 2010).

However, if high granularity is not needed by downstream analysis, then the use of EQ syntax might be time inefficient. For example, the ontology-informed phylogenetic method for reconstructing ancestral anatomies PARAMO (Tarasov et al., 2019) uses an input where a character statement is only tagged with one or few ontology term(s) (i.e., URI(s): uniform resource identifier). To date, there is no software that would facilitate this “light” version of phenotypic annotation, at the same time, doing this manually is laborious due to enormous amount of terms contained in any ontology.

Unlike its popularity in the phylogenetics community, R environment (R Core Team, 2020) is not broadly used for developing ontology-oriented software. This hinders creation of workflows that seek to integrate ontological and phylogenetic approaches within the same programming environment and, hence, inhibits development of new computational methods at the interference of the two fields (Tarasov, 2019). To fill up these gaps, we have created an open source R package `ontoFAST` that provides an interactive interface for “light” phenotypic annotation of characters with biological ontologies. Additionally, it provides functions for visualizing hierarchies of characters and ontologies using the `sunburstR` package (Bostock et al., 2020) and Cytoscape (https://cytoscape.org). Finally, `ontoFAST` provides a means to construct queries of characters against ontologies for getting a new insight into their phenotype-phenotype relationships. In turn, this enhances interoperability between ontology-oriented applications and, hopefully, will stimulate further development of ontological tools in R.

## 2 MATERIALS AND METHODS

### 2.1 ontoFAST Availability

The current stable version of the package `ontoFAST` requires R 3.5.0 and is distributed under the GPL license. The package can be downloaded from CRAN at https://cran.r-project.org/web/packages/ontoFAST/index.html, its development version is available at https://github.com/sergeitarasov/ontoFAST. The detail tutorial is given at https://github.com/sergeitarasov/ontoFAST/wiki.

### 2.2 Implementation of ontoFAST

`ontoFAST` is developed using `Shiny` (RStudio, Inc, 2020) that enables building interactive web applications straight from `R`. The interactive interface of `ontoFAST` can be run either from within `RStudio` (RStudio Team, 2021) or any web browser. `OntoFAST` uses functions from `ontologyIndex` package (Greene et al., 2017) for parsing and manipulating ontologies. It also depends on `visNetwork` package (Almende B.V. et al., 2019) for interactive visualization of ontology graphs.

### 2.3 Data

We tested `ontoFAST` by using it to annotate two character matrices. All these datasets are included in the package and tutorial. One matrix with 392 characters (dataset Sharkey_2011) from the large-scale Hymenoptera (sawflies, wasps, ants and bees) phylogeny (`Sharkey` et al., 2012) was annotated using the Hymenoptera Anatomy Ontology (dataset `HAO`) (Yoder et al., 2010); the annotations are stored in `Sharkey`_2011_annot dataset.

Another matrix of 232 characters from the dung beetle (Coleoptera: Scarabaeinae) phylogeny (Tarasov, 2017) was annotated using dung beetle ontology (`Scarab dataset`), the annotations are stored in `Tarasov_2017_annot`. The Scarab ontology was developed from `HAO` by enriching it with anatomical terms specific for dung beetle since beetles, so far, lack any comprehensive anatomy ontology. `Scarab` is an informal and experimental ontology and should be used with caution in other studies.

## 3 RESULTS AND DISCUSSION

### 3.1 Input and output data

The character annotation using `ontoFAST` requires two initial pieces of data: a list of character statements and a biomedical ontology. The list of character statements can be imported into R as a csv table or a vector of text strings. To import statements from a character matrix stored in the widely-used NEXUS format, one can open it in a popular software Mesquite (Maddison and Maddison, 2018) and copy character statements into any software that supports csv format (e.g. Microsoft Excel or Atom https://atom.io/).

Any organism-specific anatomy ontology or supporting ontologies [e.g. PATO, BSPO (Dahdul et al., 2014), RO (Mungall et al., 2021)] can be used for annotating characters in `ontoFAST`; the selected ontology should be in OBO format and can be read with get_OBO() function from `ontologyIndex` package.

~~~
install.packages(“ontoFAST”)
# install.packages(“igraph”)
library(“ontoFAST”)
hao_obo<-get_OBO(system.file(“data_onto”, “HAO.obo”, package = “ontoFAST”),
extract_tags=“everything”, propagate_relationships = c(“BFO:0000050”, “is_a”))
data(Sharkey_2011)
~~~

The output of `ontoFAST` is a list object that contains annotations: character IDs and associated URIs (or names) of ontology terms. This list can be used for queries, exported to csv format, or, using special functions provided by ontoFAST, exported to third-party applications.

### 3.2 Annotating characters using ontoFAST

Prior to running the interactive interface, the read-in data have to be preprocessed in R console using the following three steps. First, run `onto_process()` function to combine ontology and character statement into a single object of ontology-index class (Greene et al., 2017); this function automatically parses synonyms from the ontology, the argument `do.annot = TRUE` runs fuzzy matching of characters against the ontology and suggests candidate terms for annotation. Second, create a new environment to store a variable that will serve as an input and output for the interactive mode; the new environment should be called ontofast (other names will not work), it enables global usage of the input variable for the functions operating during the interactive session. Third, use the function `make_shiny_in()` to create an object in the ontofast environment; the name of this variable is taken as an argument by `runOntoFast()` to launch the interactive session.

~~~
hao_obo<-onto_process(hao_obo, Sharkey_2011[,1], do.annot = FALSE)
ontofast <-new.env(parent = emptyenv())
# creating shiny_in variable to serve as an input and output for runOntoFast()
ontofast$shiny_in <-make_shiny_in(hao_obo)
# running the interactive session
runOntoFast(is_a = c(“is_a”), part_of = c(“BFO:0000050”), shiny_in=“shiny_in”,
file2save = “OntoFAST_shiny_in.RData”)
~~~

The interactive interface consists of four panels (Fig. 1). The **ontology panel** shows an interactive graph where nodes are the ontology classes and edges are usually *part_of* and *is_a* relationships. The **customize panel** on the top of the window allows selecting relationships to display and navigate to a required term by typing a few letters of its name. The **information panel** shows ID, synonyms, and definition of the term selected in the navigation panel.

**FIGURE 1.**
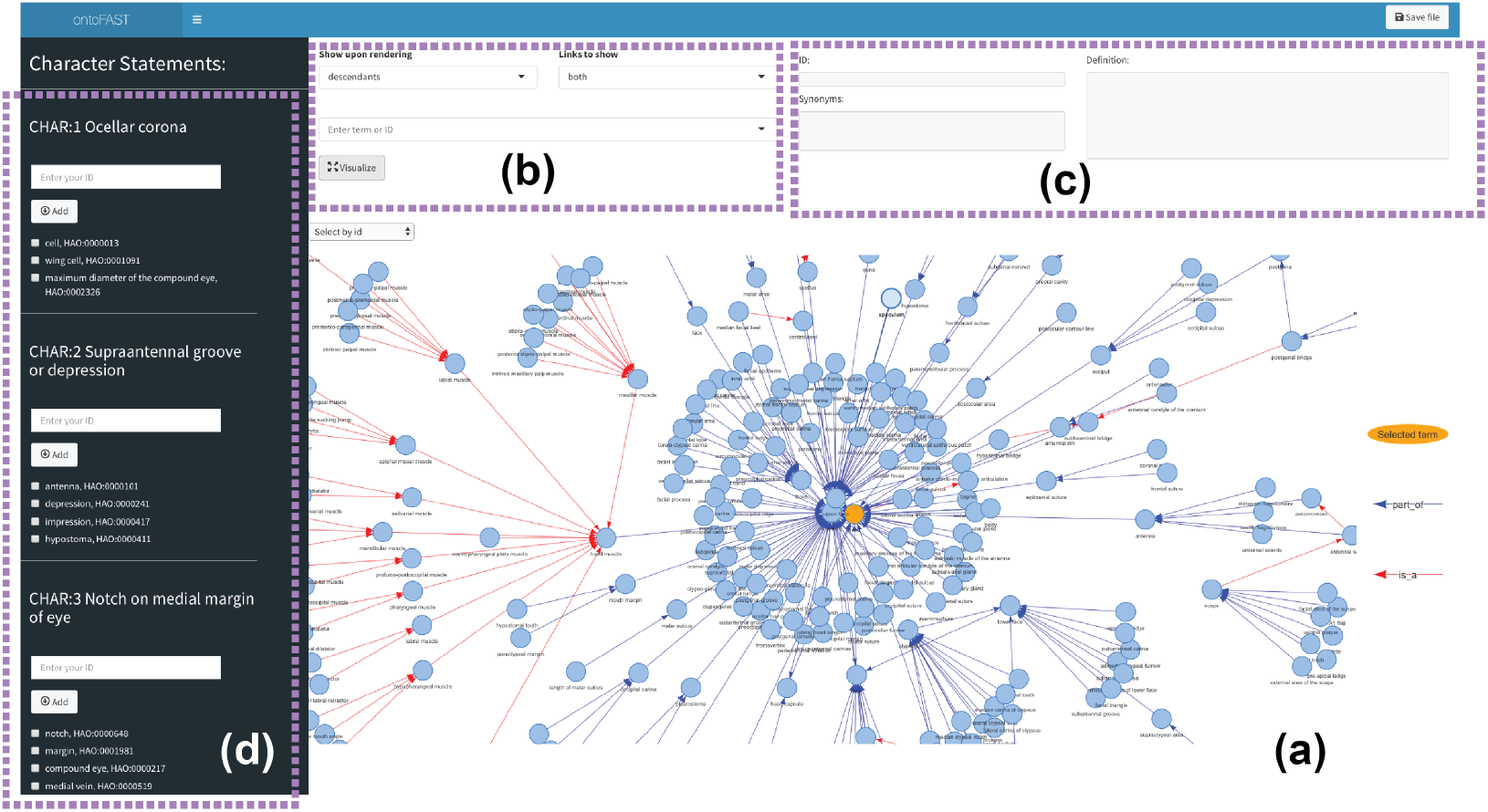
The interface of ontoFAST. **(a)** Ontology panel. **(b)** Customize panel. **(c)** Information panel. **(d)** Character panel.

The leftmost **character panel** shows the character statements. There are three ways to annotate them: (1) if you ran fuzzy matching with `onto_process()` the candidate terms are shown below the “Add” button and can be selected by checking the respective box(es); (2) click on a node in the **ontology panel**, move the cursor to the **character panel** and click the respective “Add” button; (3) paste term URI right in the **character panel**. Every character can be annotated with more than one term.

Upon the annotation is complete you can close the window and return to the console mode. The characters and their annotation are stored in the lists `ontofast$shiny_in$terms_selected and ontofast$shiny_in$terms_selected_id`. Consider saving your data, by clicking the “Save file” button in the top right corner, while in the interactive mode. This will help to avoid risk of loosing annotations, if R session crashes; ontoFAST saves data to the file specified via `file2save` argument in `runOntoFast()`. The created annotations can be further used in downstream analyses or saved as csv files.

~~~
out <-list2edges(ontofast$shiny_in$terms_selected_id)
write.csv(out, “annotations.csv”)
~~~

### 3.3 Visualizing and queering annotations

#### Visualizing with sunburstR and Cytoscape

Having characters linked with ontology may provide new insight into their relationships. We use the annotations produced with `ontoFAST` for Hymenoptera and dung beetles (see the Data section) to demonstrate it. The hierarchical structure can be visualized using sunburst plot from the sunburstR package (Bostock et al., 2020). This plot shows relational hierarchy using a series of rings; each ring corresponds to a level in the ontological hierarchy – the inner circles represent ontology classes and outermost circle represents the annotated characters (Fig. 2b-d). The function `paths_sunburst()` automatically convert `ontoFAST` data to sunburstR format.

**FIGURE 2.**
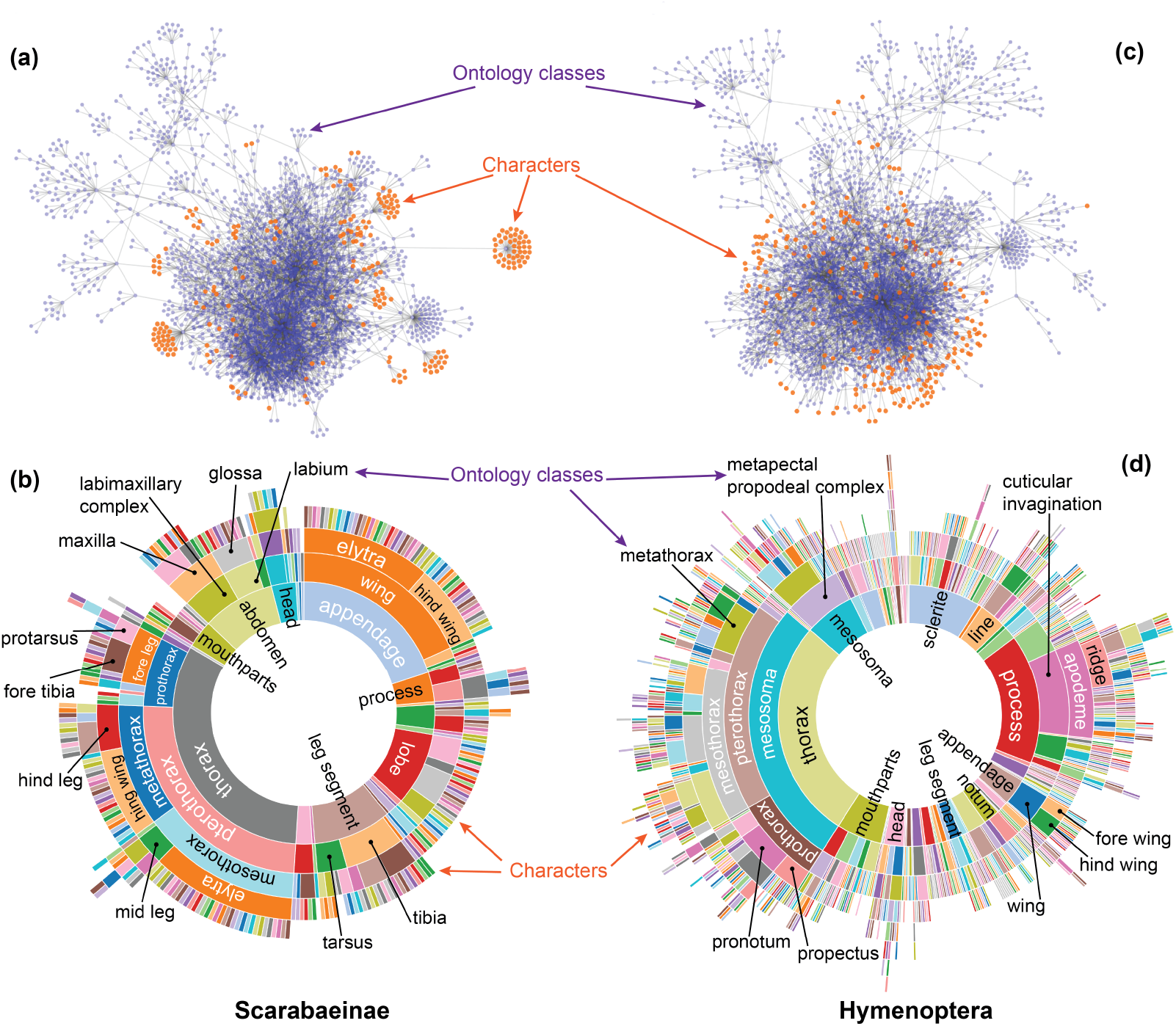
Visualization of characters annotated using `ontoFAST` for Hymenoptera and Scarabaeinae (dung beetles). **(a-b)** The network of ontology classes and the linked characters produced using Cytoscape. **(b-d)** The plots show hierarchy of characters and ontological classes, produced using sunburstR package.

The annotations and ontologies can be also exported to `Cytoscape` using `export_cytoscape()` function for further manipulation and visualization (Fig. 2a-b). Cytoscape is an open source software for visualizing complex networks and integrating them with any type of attribute data.

#### Querying

Our package has a set of functions for running simple queries with the annotated characters. The function `chars_per_term()` calculates the number of characters for each ontology class; `get_ancestors_chars()` return all shared ancestral ontology classes for a set of characters; and, `get_descendants_chars()` returns all characters descending from a given ontology class.

## 4 AUTHORS’ CONTRIBUTIONS

ST conceived and designed the package; IM and ST annotated Hymenoptera and Scarabaeinae datasets; all authors contributed to designing presented annotation procedure and wrote the paper.

## 5 DATA ACCESSIBILITY

The code of `ontoFAST` version xxx including all data will be archived on Zenodo upon acceptance.

